# Spatial phylogenetics of butterflies in relation to environmental drivers and angiosperm diversity across North America

**DOI:** 10.1101/2020.07.22.216119

**Authors:** Chandra Earl, Michael W. Belitz, Shawn W. Laffan, Vijay Barve, Narayani Barve, Douglas E. Soltis, Julie M. Allen, Pamela S. Soltis, Brent D. Mishler, Akito Y. Kawahara, Robert Guralnick

## Abstract

Broad-scale quantitative assessments of biodiversity and the factors shaping it remain particularly poorly explored in insects. Here, we undertook a spatial phylogenetic analysis of North American butterflies via assembly of a time-calibrated phylogeny of the region coupled with a unique, complete range assessment for ~75% of the known species. We utilized a suite of phylodiversity metrics and associated environmental data to test whether climate stability and temperature gradients have shaped North American butterfly phylogenetic diversity and endemism. We also undertook the first direct, quantitative comparisons of spatial phylogenetic patterns between butterflies and flowering plants in North America. We expected concordance between butterflies and angiosperms based on both shared historical environmental drivers and presumed strong butterfly-host plant specializations. We instead found that biodiversity patterns in butterflies are strikingly different from flowering plants in some regions of the continent. In particular, the warm desert regions of the southwestern United States and Mexico showed surprisingly high butterfly phylogenetic diversity and endemism, in contrast to much lower values for angiosperms. Butterflies did not show patterns of phylogenetic clustering as found in flowering plants, suggesting differences in habitat conservation between the two groups. Finally, we found weak relationships and spatially structured biases in relative branching timing between angiosperms and butterflies. These results suggest that shared biogeographic histories and trophic associations do not necessarily assure similar diversity outcomes. The work has applied value in conservation planning, documenting warm deserts as an important North American butterfly biodiversity hotspot.

## Introduction

Insect biodiversity patterns are very poorly understood across broad spatial, temporal, and phylogenetic scales. These shortfalls stand in stark contrast to our knowledge of vertebrate and flowering plant biodiversity, especially in North America, where rapid efforts to close phylogenetic and spatial information gaps^1–3^ have provided novel insights into the shorter and longer-term processes structuring biodiversity and how best to preserve natural heritage for the future^4^. One approach to quickly expand the knowledge base of insect biodiversity, and preserve it in the face of accelerating terrestrial declines^5^, lies in focusing on clades where existing data are already dense, but not yet fully integrated. Such efforts also provide a unique basis for direct, empirical comparisons with other lineages, such as various lineages of green plants, that are known to have strong evolutionary and ecological associations with herbivorous insects^6^.

Butterflies (Papilionoidea) serve as an ideal study group for researchers and naturalists due to their diurnal activity, often vibrant and showy colors, and specialized larval host plant associations^7,8^. Not only are they the most collected and photographed insects^9^, but many species and clades of butterflies have become models for studying diverse ecological and evolutionary processes, such as Batesian and Müllerian mimicry (e.g., butterflies in the genus *Heliconius*^10–13^), genetics and migration (e.g., butterflies in the genus *Danaus*), and adaptation to agricultural systems (e.g., the common and widespread cabbage white, *Pieris rapae*^14^). Butterflies also serve as pollinators and bioindicators of change and are one of the few insect groups where conservation agencies such as the IUCN have made at least initial assessments of species endangered status^15^.

Due to the interest of both professionals and amateurs, the natural history of North American butterflies is relatively well known with rich distributional and genetic data resources readily available. There are approximately 1900 species of butterflies in North America^16^, and natural history and genetic data exist for nearly 1500 species of them. This abundance of data positions butterflies as one of the best insect groups for asking broad-scale questions about the structure and drivers of diversity. These strong data sources enable moving beyond simple taxic summaries of diversity, such as species richness, and towards a more comprehensive, process-oriented understanding of how diversity is evolutionarily structured at the continental scale. Despite such potential, a synthetic, broad-scale phylodiversity analysis of butterflies (or any other insect group) and the drivers of that diversity has yet to be conducted. Even North America-wide summaries of butterfly taxic diversity have been limited^17–19^.

Butterflies are one of the most sensitive insects to changes in climate^20^. A fundamental question is how current climate and historical changes in temperature and landscape across North America have shaped butterfly phylogenetic diversity and endemism. North America is characterized by a wide range of ecosystems, a dynamic geological history, and significant insect diversity^21,22^. Butterflies are distributed across 14 broad ecoregions, ranging from the Eastern Temperate Forests to tundra and taiga in Northern Canada, tropical wet forests in southern Mexico, and the warm and cold deserts of the Southwest^16^. Landscapes across the continent have dramatically changed during the Quaternary, especially in the west, due to long-term aridification and orogeny leading to formation of the Sierra Nevada mountains, and across northern portions of the continent through cyclic patterns of glaciations^23^.

Butterflies also rely heavily on flowering plants, as sources for both adult nectar and larval food^24^. A key question is whether butterflies and angiosperms show concordant broad biogeographic patterns given these strong ecological associations and shared historical landscape and climate drivers. Recent efforts to document North American plant phylodiversity^25^ provide a data basis for direct, quantitative comparisons of butterflies with angiosperms. That recent analysis is the most comprehensive yet attempted, covering more than 19,500 plant species (out of more than 44,000 total species), found across the continent. The work presented here is the first study to directly compare spatial patterns and drivers of phylogenetic diversity between any group of insects and flowering plants at a continental scale.

Here we assembled and analyzed butterfly spatial phylogenetic diversity across North America and examined its connection to historical climate and flowering plant phylodiversity patterns. Phylogenetic approaches have two key advantages compared to traditional taxic approaches. First, phylodiversity metrics reduce reliance on species definitions; rather, branch lengths are used to calculate diversity metrics. Second, spatial phylogenetic approaches bring in evolutionary history and allow hypothesis testing, making it possible to assess, for example, whether communities are more distantly or closely related to each other than expected by chance.

We applied a set of spatial phylogenetic methods and metrics, including phylogenetic diversity (PD),^27^ phylogenetic endemism (PE)^28^ and relative phylogenetic diversity and endemism (RPD and RPE).^29^ We also employed CANAPE, which can differentiate between types of endemism found in a region,^29^ namely between recent radiations leading to neoendemism and relictual endemism leading to range-restricted groups that were once more widespread, i.e. paleoendemism. While these metrics are now commonly applied, we provide a short summary of those used here in Table 1.

**Table 1:** Summary of phylodiversity metrics and tests used.

|  |  |
| --- | --- |
| Phylogenetic Diversity (PD) | Measured as the sum of branch lengths connecting the terminal taxa present in each location (usually to the root of the tree) |
| Phylogenetic Endemism (PE) | Like PD but measured on a tree where the branches are weighted by the inverse of their geographic range (a Range-Weighted Tree) |
| Relative Phylogenetic Diversity (RPD) and Relative Phylogenetic Endemism (RPE) | Ratio of PD or PE measured on the original tree to PD or PE measured using a comparison tree with the same topology, but where each branch is adjusted to be of equal length. |
| Categorical Analysis of Neo- and Paleo-Endemism (CANAPE) | Geographic centers of endemism are identified, first as being significantly high in either the numerator or the denominator of RPE (or both), and then classified as paleo, neo or mixed based on whether the RPE ratio is significantly high or low. |
| Randomization Tests | These metrics are tested for statistical significance using a spatially structured randomization that re-assigns terminal taxon occurrences on the map, subject to two constraints: the range size of each taxon and the richness of each locations are held constant, |

We used these metrics of phylogenetic diversity and endemism to test hypotheses about a set of potential drivers and associations, including a unique, direct empirical comparison between butterflies and flowering plants. These same techniques also document centers of diversity and endemism that may differ from plant or vertebrate groups and inform conservation prioritization. Based on a recent analysis of North American plant phylodiversity, we made the following predictions:

1. Regions that are warmer and have remained more stable over time will have higher phylodiversity (PD)^30,31^. Stable areas, whether warm or cold, should have significantly higher than expected PD because they have had the most time to accumulate lineages^32^, along with specializations that may structure communities to avoid competition^33^.
2. Relative phylogenetic diversity (RPD) will be higher than expected in areas that have been most stable, accumulating more long-surviving, older lineages^33^. In North America this includes the eastern and southernmost portions of the continent, as seen in the results from flowering plants. Areas with high topographic heterogeneity and that have been most climatically unstable, such as recently deglaciated areas in the north and portions of the west, will have significantly lower than expected RPD.
3. Butterfly phylogenetic endemism in North America will align with hotspots of high angiosperm endemism. Hotspots of neo-endemism are more likely in younger areas with higher topographic relief while areas of paleo-endemism will be highest in areas where climate and landscapes have been more stable.
4. Continental scale flowering plant and butterfly phylodiversity patterns will be highly congruent, due to co-evolutionary dynamics between the two groups and to similarities in underlying landscape and climate drivers.

## Materials and Methods

### Species Name Assembly

We consolidated a list of all North American butterfly species with their current valid names and known synonyms. We defined North America as including Canada, Mexico, and the United States, but excluded species that were endemic to islands near the North American landmass (e.g., the Caribbean), as these islands were not well included in field guides. Valid names were derived from a global checklist^34^ and were augmented via assembly of synonymies from *Lepidoptera and other life forms* database (Funet^35^) and Wikipedia^36^ using R package taxotools. The augmented master list was used to normalize names from resources that contained expert-assessed maps and assembled names used in field guide resources^7,37^. Once names were normalized to a consistent, accepted name, we used those names and associated synonyms to: (1) re-assign normalized names to those digitized species range maps (see below for range map assembly) where normalization was required and (2) search GenBank and other key resources for matching genetic data, to construct a North America-specific butterfly phylogeny.

### Range Maps and Digitization

Range maps were digitized from field guides covering the USA and Canada^7^and Mexico^37^ for each species included in our species list. Digitizing steps included generating high-resolution scans, georeferencing the resulting images, and then manually tracing polygons based on those scans in QGIS version 3.2^38^. Less than 1% of the species in field guides did not have an associated range map; in those cases we used occurrence records and descriptions in field guides to estimate the ranges. Range maps were combined into a single shapefile consisting of many spatial polygons which were clipped to only terrestrial areas within North America. We provide details about range map digitization and a rigorous approach to quality control of maps in Supplemental Information 1.

Many butterfly species have ranges that extend beyond the borders of North America. Calculations that involve range-weighting, such as phyloendemism, should ideally rely on globally complete phylogenies and range estimates. Here, we partially compensated for this currently unattainable goal by determining a coarse estimate of overall range extents using country-level range maps for every species in our list where needed. We generated these country-level range maps utilizing three separate resources: (1) Country-level ranges from Funet^35^; (2) GBIF data for all relevant species and extracting country-level data from these records; (3) Data from a trait database that was assembled from field guides and other published sources^39^. Supplemental Information 1 describes more details on country list production and quality control.

A 100 km by 100 km resolution grid at the global scale was projected to a North America Albers equal area conic coordinate system. A species was considered present in the cell if the cell centroid was within the species’ range map or if the distance from each centroid to the nearest edge of the species’ range map was less than 1 km.

### Sequence Data Acquisition and Dataset Construction

We compiled sequence data for 13 common markers (1 mitochondrial and 12 nuclear genes) used in butterfly phylogenetics (see Supplemental Table 1). These sequences were obtained from GenBank^40^, Barcode of Life Data System (BOLD)^41^, and by extracting tissues and sequencing with Sanger and target capture sequencing for species that lacked genetic data. We accumulated existing marker data from GenBank using a python toolkit (https://github.com/sunray1/GeneDumper) developed by the lead author. This toolkit automatically fetches sequences from GenBank and uses a rigorous, automated cleaning workflow to choose the best matching sequences for further processing. We also queried for relevant loci and sequences in BOLD^41^, as some of these data records are not reflected in GenBank. BOLD data were assembled using its API (http://www.boldsystems.org/index.php/resources/api). Target capture sequencing utilized Anchored Hybrid Enrichment (AHE)^42^ on 224 butterflies with the Butterfly 1.0^43^ and 2.0^44^ target capture sets. Both of these kits include the 13 markers of interest. We also generated new mitochondrial cytochrome C oxidase subunit I gene (COI) sequences by extracting DNA from dried pinned museum specimens in the Florida Museum of Natural History, McGuire Center for Lepidoptera and Biodiversity at the University of Florida. These tissues were shipped to the Canadian Centre for DNA Barcoding (CCDB; Guelph, Canada) for sequencing of the standard 658-bp region of COI. See Supplemental Information 2 for additional information on sequence acquisition. We created FASTA files for each locus which were aligned using MAFFT v.7.294b^45^ and concatenated into a single alignment with FASconCAT-G v.1.02^46^. The concatenated alignment was 12,361 bp in length and had 69% missing data. Most species (99%) were represented by COI. Without COI, missing data increased to 76% across the remaining 12 loci.

### Phylogeny Construction

All loci included in this study were protein-coding, and therefore the alignment was partitioned by gene and codon position, resulting in 39 partitions. PartitionFinder v.2.1^47^ was used to choose the best partitioning scheme and nucleotide substitution models. A phylogenetic analysis of North American butterflies was conducted in RAxML v.8.2.10^48^ with the concatenated alignment and best partitioning scheme. We conducted 100 ML tree-searches with different random seeds, and the tree with the best log likelihood score was chosen as the final tree. We determined branch support by running 200 parametric bootstrap replicates in RAxML under the GTR+Γ+I model and a gradual “transfer” distance method implemented in BOOSTER^49^. A final check was made to ensure that tip names were consistent with the species names from range products.

The interpretation of phylodiversity metrics depends on the units of the branch lengths in the phylogeny, e.g., the amount of “feature diversity” contained in a region when using a phylogram or the amount of “evolutionary history” in a region when using a chronogram^50^. We focus here on chronograms, with their explicit focus on the age of lineages and communities and therefore produced a time-calibrated tree. Divergence times were calculated using penalized likelihood in TreePL^51,52^ following a congruification approach^53^. In particular, we obtained node calibrations from Espeland et al.^43^, extracting date ranges from each of the six family nodes that were concordant with our phylogeny and using those as estimates in TreePL (see Supplemental Information 3 for more phylogeny reconstruction details).

### Analysis of Phylogenetic Diversity and Endemism

The spatial data set and the phylogeny described above were imported into Biodiverse v.3.0^54^. Tips on the tree were mapped to species in the spatial data set to calculate species richness (SR), phylogenetic diversity (PD), and phylogenetic endemism (PE) metrics for equal-area square grid cells (100 × 100 km). The imported mapping extent was global (as described above) in order to calculate range size metrics, but our analysis region for phylodiversity metrics were constrained to North American grid cells using spatial constraints in Biodiverse. This approach ensured that PE metrics take into account overall range sizes of the terminals in the tree, including their extent outside of the continent. *Relatives* of the terminals that occur elsewhere in the world are not included, but this is the best approach possible to estimating PE until global analyses are feasible.

We also calculated relative phylogenetic diversity (RPD) and relative phylogenetic endemism (RPE)^29^. These are ratios of PD and PE on the original tree compared to a phylogeny with the same topology but with equal branch lengths. These metrics can provide useful information about areas with concentrations of significantly longer or shorter than expected phylogenetic branches. For example, areas that have more recently radiated taxa are expected to have lower chronogram-derived RPD. All cell-based values for PD, PE, RPD, and RPE were exported from Biodiverse for mapping and further analysis.

### Randomization Tests

Phylogenetic diversity and endemism measurements are expected to be highly correlated with taxic diversity (species richness), because each taxon added to a community must also add to the overall PD. If the co-occurring taxa are randomly distributed on the tree, the correlation should be tight, so a key step forward is to move beyond simply reporting summary measures, and test whether phylodiversity values are higher or lower than expected compared to null models of randomized communities^55^. This is achieved through a randomization approach, where species occurrences within North America are randomly reassigned to grid cells while holding constant the richness of each cell and the range size of each species. Values for PD, PE, RPD, and RPE were then calculated for each randomization iteration, creating a null distribution for each grid cell. A two-tailed test was then applied to the PD, PE, and RPD randomizations to determine whether the observed values were significantly high or low when compared to the null distributions. We utilized a 40-core Dell Xeon PowerEdge standalone server to parallelize creation of 500 random realizations per grid cell across all cores. Only observations within the North American study region were randomized, with the outside regions held constant. Output randomized results were merged, exported as GeoTIFF grids and re-imported in R for downstream analysis.

RPE randomizations enable a means to categorize different types of phylogenetic endemism. This method, called *Categorical Analysis of Neo- And Paleo-Endemism* (CANAPE^29^), is a two-step approach that first selects grid cells that are significantly high (one-tailed test) in either the numerator or the denominator of RPE, then uses a two-tailed test of the RPE ratio to determine four possible outcomes per cell: higher than expected concentrations of range-restricted short branches (i.e. neo-endemics); long branches (i.e., paleo-endemics); a mixture of both types; or no significant endemism. Endemism measures, including randomizations, were calculated in Biodiverse and the categorization method for CANAPE was run in R v.3.6.3^56^ to determine per-grid-cell phylogenetic endemism types, and to plot those results spatially.

### Drivers of Phylodiversity and Endemism

#### Assembly of explanatory variables for diversity and endemism patterns

We used seven variables to analyze the observed phylogenetic diversity and endemism patterns. These included four bioclimatic variables (annual mean temperature, annual precipitation, temperature seasonality [standard deviation * 100], and precipitation seasonality [coefficient of variation]^57^), two climate stability variables (temperature stability and precipitation stability), and elevation. The climate stability variables represent the inverse of the mean standard deviation between equally-spaced 1000-year time slices over the past 21,000 years and were provisioned from Owens and Guralnick^58^. These seven layers were chosen because they capture the geographic variation in climate stability over a significant transition from a full glacial to interglacial time-period, which likely is representative of similar transitions that occurred repeatedly in the Pleistocene^59^. Elevation values at this scale also provide a reasonable proxy of current topographic heterogeneity. All environmental variables were scaled to a mean of zero and SD of one and resampled to 100 km × 100 km.

#### Testing Importance of Climate and Topography Drivers

Four diversity metrics (PD, RPD, and randomization tests for both measures) were utilized to test the most predictive explanatory variables of diversity and diversity significance. Phylogenetic significance analyses were derived from the randomized metrics and divided into binomial datasets, where significantly high values were scored with the value one and significantly low values assigned zero. Cells with non-significant values were excluded from the binomial regressions.

We fit generalized linear models using climatic variables described above as predictors and phylogenetic metrics as response variables. For the PD and RPD analyses, we used the Gaussian distribution; for models examining PD/RPD significance we used the binomial distribution. We used the dredge function from the package MuMIn^60^ in R version 3.6.2 to examine all possible models. We used an information-theoretic approach using Akaike’s Information Criterion (AIC) to rank models^61^. Models that were a subset of another model examined were not considered to be competitive if within delta AICc ≤ 2. We examined the collinearity of variables of the models by calculating variance-inflation factors (VIF) using the car package, and models with VIF ≥ 5 were also not considered as competitive models. We used delta AIC values and Akaike weights (wi) to rank competing models.

#### Comparison of Butterfly and Plant Phylodiversity

We re-ran a recently published analysis of seed plant phylodiversity^25^ but excluded gymnosperms in order to compare our results to spatial phylogenetics patterns for North American flowering plants. This re-analysis used the same methods as in Mishler et al.^25^ and resulted in inclusion of 19,173 angiosperm terminal taxa (and exclusion of 476 gymnosperms) across North America. We applied the same metrics for angiosperms as for butterflies, and used nearly the same spatial extent, only excluding a small portion of the southern tip of Mexico, at 50-km resolution. We resampled gridded analysis products from that study into the same 100-km resolution as the butterfly grids using bilinear interpolation for comparisons of associations between butterfly and plant phylodiversity metrics. We next generated univariate linear regression models, where plant PD, RPD, and PE values were used as the predictor variables and the corresponding butterfly PD, RPD, and PE values were the response variables. The residuals of these models were then mapped spatially to display where plant diversity metrics under- or over-predicted butterfly diversity. Comparison of PD, RPD, and CANAPE significance for butterflies and flowering plants was done by visual inspection because a summary test was gauged to be superfluous given the striking regional differences between the two groups.

## Results

### A Phylogeny for North American Butterflies

The ML analysis resulted in a tree of 1,437 North American butterfly species (Supplemental Figure 1). A total of 1,437 (74.6%) known butterfly species had sequence data already available or were *de-novo* sequenced for COI. *De-novo* sequencing led to the addition of 140 species lacking data in existing repositories. Of these 140 species, 96 species (68.6%) were distributed only in Mexico. Although the backbone of the butterfly tree was constrained at the family level, subfamily and tribe-level relationships generally agree with those of prior studies, (Supplemental Information 3) and clade-based ages were largely congruent with recent butterfly-wide dating analyses (Supplemental Table 2). We recovered a median bootstrap value across the entire tree of 86 using transfer bootstrap expectation^49^.

### Observed Patterns of Diversity and Endemism

Maps of observed species richness (Figure 1A) and phylogenetic diversity (Figure 1B) both documented highest diversity primarily in the tropical dry and wet forests in Mexico and the lowest values across the arctic of Canada. Patterns of richness and PD are complex in western North America, likely reflecting the the heterogeneous landscape, with peaks in areas adjoining the Sierras and Rocky Mountains. Relative phylogenetic diversity (RPD) peaked in wet and dry tropical forests in Mexico, and remained uniformly high across the Eastern Temperate Forest, Great Plains, and southern deserts (Figure 1C). By comparison, observed RPD was lower across much of the temperate Intermountain West, the Mediterranean regions of California, and into northern ecosystems such as boreal forests and taiga. Phylogenetic endemism (PE; Figure 1D) showed the same general latitudinal gradient as PD and RPD, but included areas of higher phylogenetic endemism along the temperate Sierra Madre mountain ranges in Mexico, and in the coast ranges in the Pacific. PE was overall higher in the temperate west than in the east and associated with transition zones in the Rockies and Sierra Nevada. However, a key limitation with the current analyses is the coarse spatial resolution, which limits finer localizations of these patterns along spatial and environmental gradients^62^.

**Figure 1.**
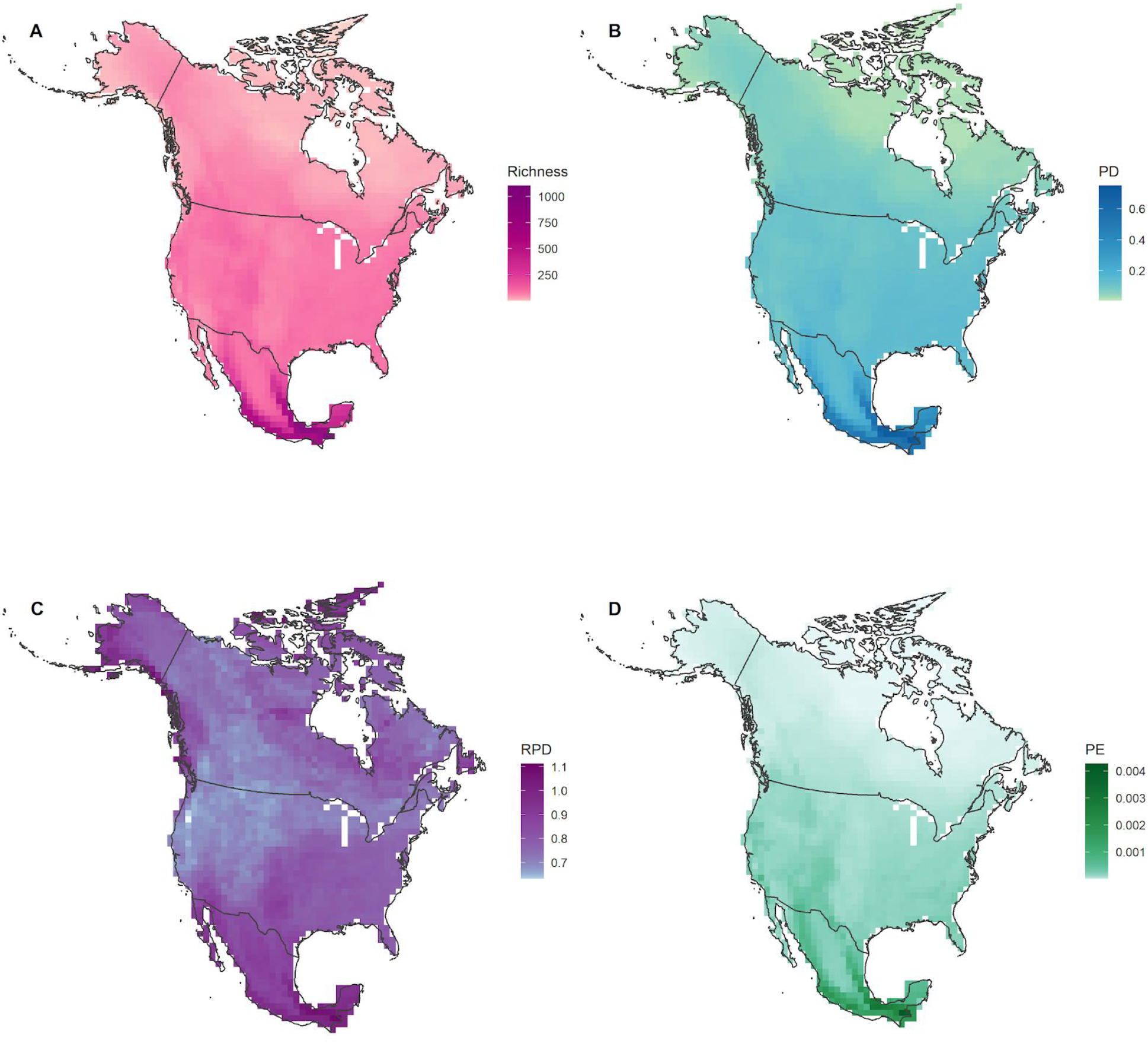
Observed values for North American butterflies for: (A) taxic richness, (B) phylogenetic diversity (PD), (C) relative phylogenetic diversity (RPD), and (D) phylogenetic endemism (PE).

### Spatial Randomization Tests

We uncovered highly regionalized patterns of overdispersion and clustering based on PD randomizations (Figure 2B). All boreal, taiga, and tundra regions showed lower than expected PD, indicative of phylogenetic clustering. Most of the temperate regions in the west, including diverse ecoregions in cold deserts, west coast forests, and Mediterranean portions of California, also displayed clustering. In contrast, tropical wet and dry forests, the most phylodiverse areas in North America, showed higher than expected PD, or phylogenetic overdispersion, when compared to null models. We also note that portions of the south-central semi-arid prairies also showed higher than expected phylodiversity. The southern, warm deserts and eastern temperate forests did not show significantly high or low PD.

**Figure 2.**
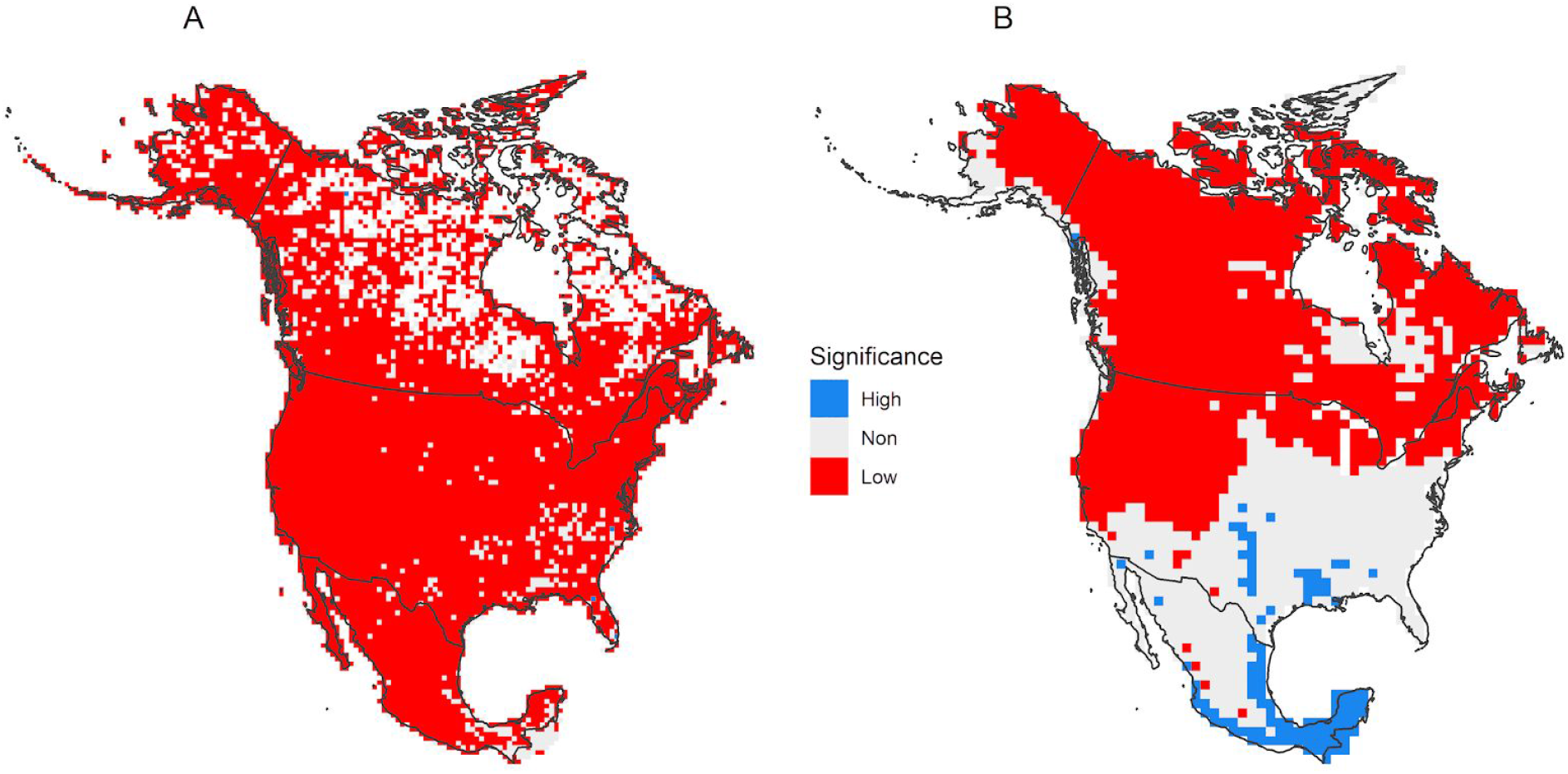
Statistical significance of phylogenetic diversity (PD) for (A) angiosperms and (B) butterflies across North America. Areas with significantly high values have taxa that are less closely related than expected by chance (blue), while areas with significantly low values have taxa that are more closely related than expected by chance (red).

The RPD randomization indicated that southern portions of North America have communities containing longer branches than expected under null models (Figure 3B). This included not only tropical regions, but also semi-arid highlands, and southern deserts into semi-arid plains and prairie. On the other hand, shorter than expected branch lengths were found across much of the Sierra Nevada, Rockies, and Intermountain West. We found no significant RPD in the Eastern Temperate Forest, northern Great Plains, and northernmost portions of North America.

**Figure 3:**
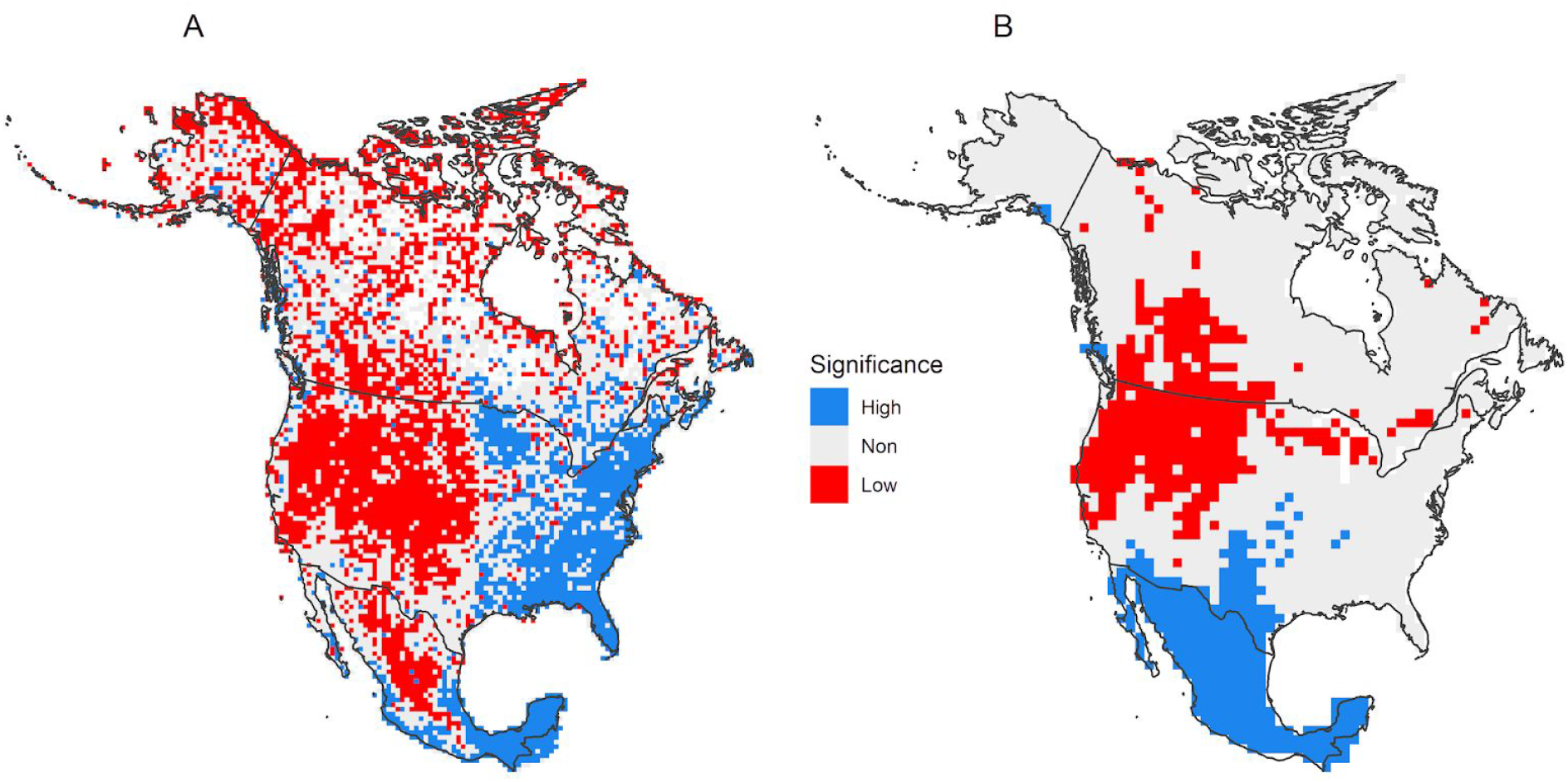
Statistical significance of relative phylogenetic diversity (RPD) for (A) angiosperms and (B) butterflies. Areas in blue have significantly longer branches than expected; areas in red have significantly shorter branches than expected.

### CANAPE

Regions of significant neoendemism were located in the California Mediterranean region and western forests, including the Cascades, Coastal Ranges, and the Sierra Nevada (Figure 4B) and in transition zones across lower-elevation regions to the East. Mixed patterns of endemism with both paleo- And neoendemics were found in predominantly warm deserts and the southeastern coastal plain and southern, subtropical portions of Florida. Sites dominated by paleoendemism were more rare, only indicated in some areas in tropical Mexico.

**Figure 4:**
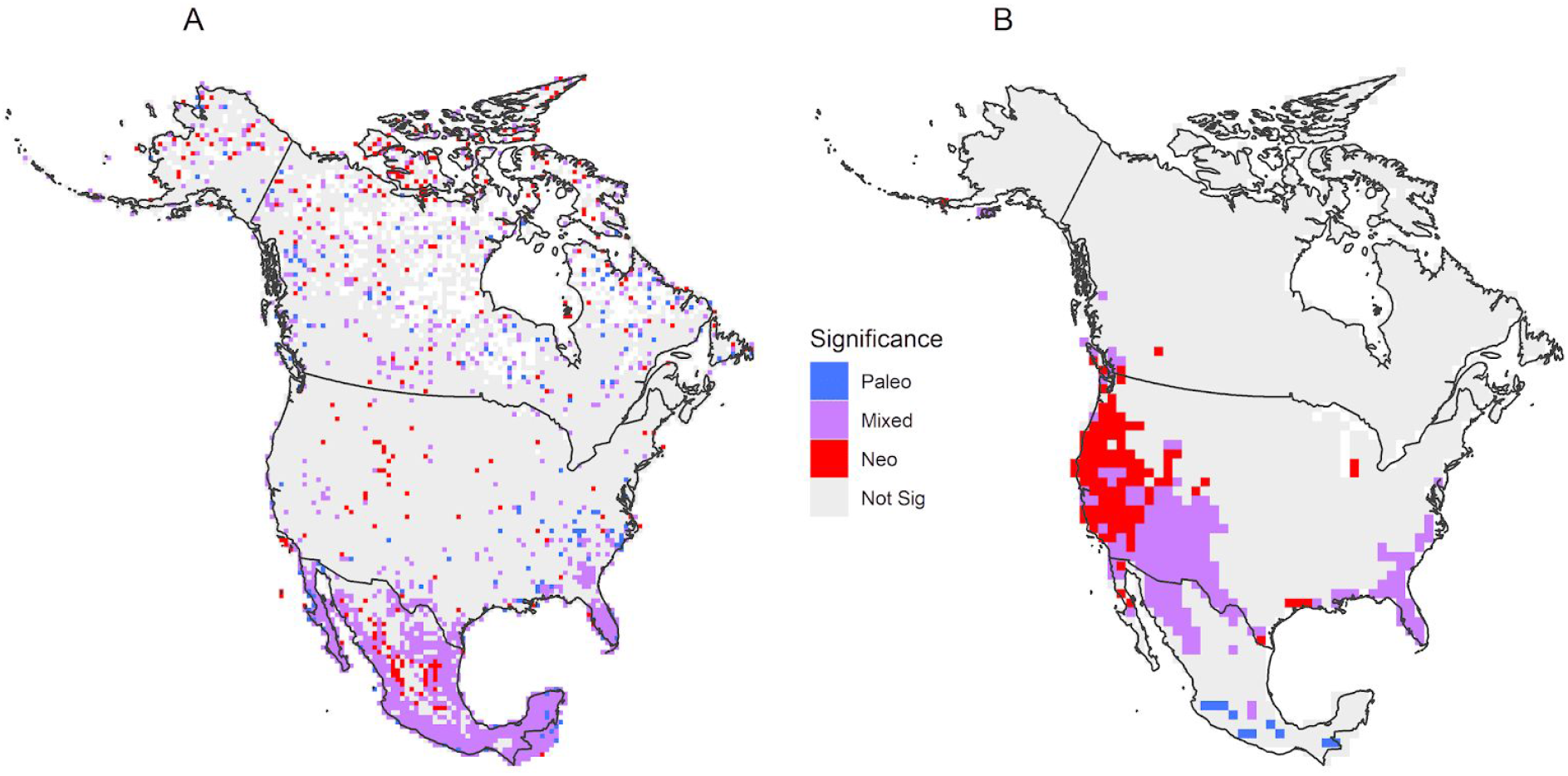
CANAPE results showing statistically significant centers of phylogenetic endemism for (A) angiosperms, and (B) butterflies. All cells that are colored have significantly high PE. Red cells have concentrations of rare short branches (neoendemism); blue cells have concentrations of rare long branches (paleoendemism), and purple cells have mixtures of neo- And paleoendemism.

### Drivers of Phylodiversity

Annual mean temperature was the most important environmental variable in predicting PD, followed by mean annual precipitation, and precipitation seasonality (Table 2). Areas that were warmer, wetter, and had more seasonal precipitation generally had the highest PD. As well, higher elevation areas and those with stable temperatures generally had higher PD, although these are weaker effects. PD significance results mirrored results for observed PD, with warmer, wetter, and more stable areas more likely to have significantly high PD (Table 2). However, elevation exhibited a pattern that was opposite that seen for observed PD, with higher areas associated with significantly lower PD.

**Table 2.** Summary of the top models for PD, RPD, significant PD, and significant RPD. Numbers in the columns indicate changes in PD, RPD, and significant PD and RPD values when variable values between locations increased by one standard deviation. Models were ranked based on the difference from the top model to the nearest competing model in Akaike’s Information Criterion (Δ AIC) and Akaike weight (Wi). Models that were a subset of another model and within Δ AIC of 2 were not considered to be competitive. Additionally, models with both temperature and temperature seasonality variables had variance inflation factors > 5, and were not considered competitive. All variables contributed significantly (P < 0.05) to models.

| Model | Temp | Prec | Temp Seas | Prec Seas | Temp Stab | Prec Stab | Elev | ΔAIC | $W_i$ |
| --- | --- | --- | --- | --- | --- | --- | --- | --- | --- |
| PD | 0.048 | 0.023 |  | 0.021 | 0.012 |  | 0.008 | -29.75 | 1.00 |
| RPD |  | 0.015 | -0.027 | 0.024 | 0.007 | 0.010 | -0.024 | -7.92 | 0.98 |
| PD Sig | 0.170 | 0.065 |  | 0.061 | 0.036 | 0.023 | -0.047 | -12.50 | 1.00 |
| RPD Sig | 0.470 |  |  | 0.064 |  | 0.040 |  | -7.81 | 0.98 |

Lower RPD in an area indicates relatively short branches, potentially indicative of more recent radiations. Results from analysis of climate and terrain drivers showed low RPD in areas that have higher temperature seasonality and higher elevation. As well, areas with low precipitation seasonality also had lower RPD. Climate stability is only a weak driver, and more stable areas show higher RPD. RPD significance tests showed that areas with significantly low RPD are in colder areas with less seasonal precipitation, and less long-term stability in precipitation regimes.

### Similarities and Differences Between Butterfly and Plant Phylodiversity

Butterflies and plants of North America displayed a similar pattern of PD (*r*^2^ = 0.34), and areas of discordance displayed moderate spatial structuring of residuals in our simple linear models with butterfly PD as a response variable to plant PD. Angiosperm PD was proportionally lower in the tropics compared to butterflies, and marginally higher than expected across especially the West Coast of North America (Figure 5). Butterflies and plants surprisingly did not have similar patterns of RPD (*r*^2^ = 0.01), and RPD showed strong spatial structuring of linear model residuals, with angiosperm RPD much higher in the west and lower in the south compared to butterfly RPD (Figure 5). Butterflies and plants both showed a pattern of having highest PE values in southern Mexico, but overall the similarity across North America was relatively weak (*r*^2^ = 0.10). The spatial residuals of the PE linear models mirrored PD in the southern portions of the continent but without spatially structured error in temperate regions of the continent (Figure 5).

**Figure 5:**
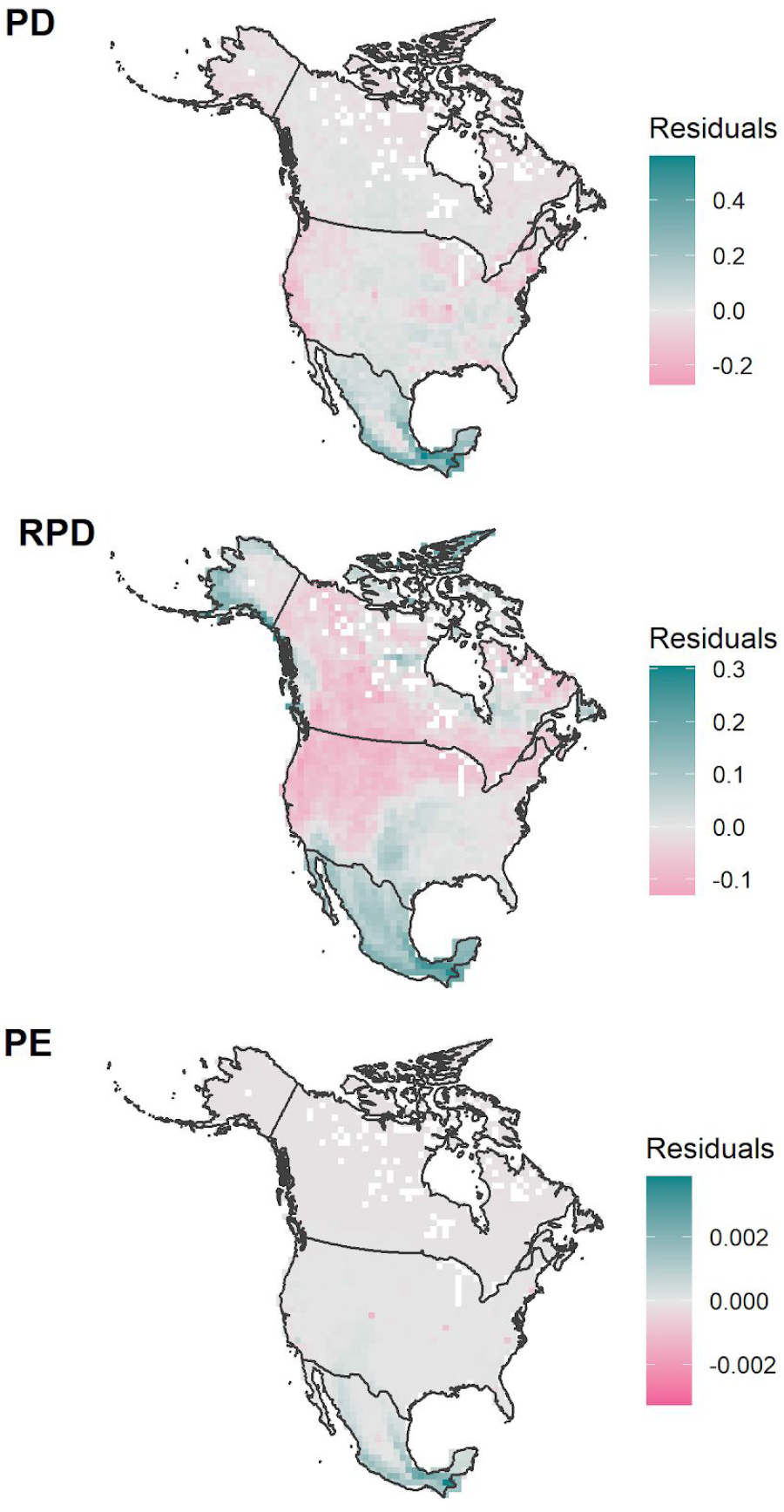
Spatial residuals of univariate linear regressions for observed PD, RPD, and PE, where angiosperm metrics were used to predict butterfly metrics. High residual values (turquoise) represent areas where butterfly values are higher than plant values.

The expectation was that drivers of flowering plant phylodiversity should directly relate to butterflies given the ecological associations between the two groups and shared biogeographic histories. However, analyses of flowering plant PD showed significantly lower than expected values across the continent, indicating phylogenetic conservatism in habitat preference and ecological filtering^25^. Butterflies showed a strikingly different pattern of PD significance, with higher than expected values in the south and lower than expected in the north (Figure 2). Angiosperms showed a strong pattern of having significantly lower than expected values of RPD in Western North America and significantly high RPD in southern Mexico and eastern North America (Figure 3). Butterflies also had significantly high areas of RPD in tropical wet and dry forests in southern Mexico, but unlike flowering plants, they also exhibited high RPD in Baja California and the American Southwest. Also unlike angiosperms, butterflies did not show significantly higher RPD in Eastern Temperate Forests. Both groups showed significantly low RPD in much of Western North America (Figure 3).

Flowering plants and butterflies showed generally discordant patterns of endemism. Centers of mixed paleo and neo-endemism for angiosperms were found in Mexico, including the Baja California peninsula, as well as in Florida and the adjoining southern coastal plain. Although CANAPE results for butterflies also showed mixed phylogenetic endemism in Florida, otherwise the results for the groups were quite different. Butterflies showed strong patterns of mixed endemism north of Mexico, in the warm deserts and portions of the colder deserts of the Southwest, along with predominantly neo-endemism in coastal regions of the West, and limited paleoendemism in southern Mexico (Figure 4).

## Discussion

We present the first continental-scale phylodiversity analyses for butterflies, focusing on North America. This analysis is notable for being relatively complete, with coarse-scale distribution data for all species, and a phylogeny with ~75% sampling of North American species. This level of completeness provides, for the first time, a well-resolved, continental-scale view of phylogenetic diversity for an entire insect suborder. We also extended the range estimates beyond North America, by gathering very coarse country-level range maps for the ~25% of butterfly species that have ranges outside of our defined North America boundaries. We argue this approach is better than simply truncating ranges of terminals for any analyses relying on a range-weighted metric, such as PE. Here, not including full ranges would have led to many neotropical butterflies having much smaller ranges that truncate at the border of Mexico rather than properly extending into Central and South America. However, even when one extends the range estimates of terminal taxa, this does not fully solve the “edge-effect” problem because the range sizes of related taxa occurring outside the study area are still being left out, which means ranges of deeper branches may be poorly estimated, affecting PE. Thus any study incorporating less than a globally complete assessment is still assessing primarily local endemism patterns.

Below we discuss how our results address key predictions regarding patterns of butterfly diversity across a continent with enormous habitat breadth, from the hot and dry deserts in the Southwest to the wet, tropical forests in the Yucatan and the cold, arid environments of the taiga and tundra. In particular, we focus on processes that are likely to have shaped this diversity, based on both examination of climatic and topographic drivers, and via comparison with angiosperms. We explicitly expected concordance of patterns and process because of strong associations between butterflies and their flowering plant hosts, and because both lineages are shaped by shared environmental changes experienced over millions of years.

### Patterns and Drivers of Phylogenetic Diversity and Endemism

Our results point especially to the importance of current climate drivers on phylogenetic diversity and endemism. We found that PD is highest in the warmest, wettest areas of the continent, along with regions of stable climates and along elevational gradients. RPD results point not to temperature but rather seasonality and precipitation as drivers. RPD was highest in regions with less temperature seasonality, but more highly seasonal rainfall patterns, perhaps representing conditions where highly divergent lineages could co-exist and where flowering plant diversity may also be unusually high. However, despite our predictions that temperature and precipitation stability would be a key predictor of PD and RPD *significance*, they were never a top predictor in any model. Rather, current climate predictors were the dominant drivers. Below we summarize these results more thoroughly by major regions across North America, focusing on synthesis across geographic distance and environmental gradients.

#### Northern North America

We define Northern North America as regions that were mostly covered in ice during the Last Glacial Maximum at ~21kya. The northern portion of the continent showed low PD, but not RPD, the former suggesting the importance of environmental filtering due to cold and seasonal conditions, and the latter suggesting possible disequilibrium from extinction-recolonization dynamics across glacial cycles. We argue environmental filtering is more likely in a volant group with high reproductive rates^63^, such as butterflies, in comparison to clades where dispersal can often lag behind changing conditions^64^. This is further supported by low PE and non-significant RPD and CANAPE results in this region (Figure 3; Figure 4), suggesting most species are wide-ranging species and not recently radiating across the North.

#### Western United States

Western areas south of past ice sheets and north of the warm deserts showed very strong patterns of significantly low PD and RPD. The former indicates potential environmental filtering given steep elevational and climatic gradients that themselves were in flux during glacial-interglacial cycling during the Pleistocene^65^ while the latter might indicate that butterflies have undergone recent radiations in these areas. The potential for recent radiations would also suggest high levels of neoendemism in the region. This was not the case for the Intermountain West and Rocky Mountains; however, we did recover a strong signal of neoendemism in Mediterranean portions of California, and western forests, extending into the Western portions of the Great Basin.

#### Southern North America

The southern portion of North America showed particularly surprising results, especially in the warm deserts. PD and RPD were both significantly high in tropical regions of North America, consistent with the tropics as a museum for butterfly diversity^66,67^. In the Temperate Sierra and warm deserts, we found significantly high RPD but no indication of clustered or overdispersed PD. This novel result suggests phylogenetically old communities of butterflies in deserts, a climate that formed recently, during the mid-Pliocene^68^. It has long been known that flowering plants found in this region are derived from related lineages in thornscrub and arid highlands that are phylogenetically much older^69^. Butterflies in warm deserts may therefore also be connected to older lineages that persisted in subtropical, yet still seasonal habitats in the southern regions, and less closely related to species in the colder deserts in the Great Basin. Bioregionalizations and their associations derived from phylogenetic beta-diversity provide a next-step means to examine such questions.

#### East of the Rockies

Areas east of the Rockies, including the Great Plains and Eastern Temperate Forests, are unremarkable in PD or RPD, which contrasts with spatial phylogenetic findings for flowering plants, as we discuss in detail below. We were particularly surprised that the Great Plains region, which became grassland-dominated during cooling in the Miocene and Pliocene, did not show accumulation of younger than expected lineages (i.e. significantly low RPD), as has been documented in other groups^25^. As well, tropical regions of Florida were not significantly higher in PD or RPD. However, tropical Florida and the nearby coastal plain do show significantly high levels of mixed PE, aligning with a known plant biodiversity hotspot^70^.

### Comparisons of Spatial Phylodiversity Patterns Between Butterflies and Flowering Plants

We expected strong concordance of spatial phylogenetic diversity in butterflies and flowering plants given strong ecological associations and shared abiotic drivers that have played out over long evolutionary timeframes^71^. This expectation is generally borne out in the Western and Northern portions of the continent, both shaped by recent perturbations including glaciation and aridification. However, our analyses also revealed striking differences. For example, butterflies and flowering plants do not show similar patterns of RPD, suggesting that diversification timings between butterflies and plants may not be associated at the scale and extent of this analysis. The reasons for these differences may be partially methodological. However, these results also suggest that shared historical forces and strong ecological associations can still lead to divergent historical and current biogeographic outcomes. We discuss more about methodological and biological rationales for similarities and differences below.

Although we used consistent PD metrics across both studies, allowing for direct comparisons of outputs, sampling completeness varies dramatically between flowering plant and butterfly analyses. Sampling of flowering plants, which encompasses more than an order of magnitude more diversity than butterflies^25^, is more incomplete in terms of both phylogenetic (ca. 44% of taxa included) and spatial distribution information. As problematic, both phylogenetic and spatial sampling is known to be biased. Some regions, especially in the North, are still nearly unsampled in terms of digitally accessible flowering plant specimen records, based on results in Mishler et al.^25^. This contrasts sharply with other regions, such as coastal California, where species sampling is mostly complete at this scale.

These differences in completeness of sampling and spatial bias make strong assessments of patterns more challenging. Mishler et al.^25^ recovered a pattern of significantly low PD across the continent for seed plants, and our more phylogenetically restricted analysis of flowering plants shows the same result. This suggests that co-occurring species are always more closely related than expected by chance compared to the full pool of species. This result, not seen in butterflies, might be affected to some extent by incomplete sampling, but it is such a strong and uniform result that it likely points to some fundamental differences in evolutionary ecology between butterflies and plants. Significantly low PD (phylogenetic clustering) is most often taken to indicate habitat filtering due to phylogenetically conserved habitat preferences which result in close relatives tending to occur together in communities. It may well be that habitat preference has a higher level of conservation in seed plants than in butterflies, a possibility in need of future research.

While it has long been known that Western North America has been dramatically reshaped by regional tectonism, orogeny, and climatic changes, the full magnitude of those impacts on flora and fauna besides vertebrates^72^ are just now starting to be understood^73^. Plant spatial phylogenetic work^25^ has confirmed this in a spectacular fashion, with Eastern Temperate Forests showing significantly older plant lineages than in the Great Plains and western portions of North America, both strongly shaped by cooling and aridification, showing more recent diversifications. We expected to find congruent results when examining RPD in butterflies. However, RPD between the two groups is not strongly correlated, and plant communities are comparatively older in the West compared to butterflies, based on spatial residual plots (Figure 5). As well, while some portions of the West show lower than expected RPD for both plants and butterflies, suggestive of more recent radiations there compared to other regions, we did not recover higher than expected butterfly RPD in the East as we did in plants.

These results suggest that less stable areas such as northern and western portions of North America may show moderate discordance when comparing across groups at different trophic levels, but in a consistent manner. Butterflies likely have diversified in the shadow of a persistent, highly diverse angiosperm-dominated forest in eastern temperate North America. Given massive inequality in numbers of butterfly to plant species in the region, providing ample opportunity for evolving new host relationships, and long-term stability, the overall effect is likely equilibrium between the two groups. In the West, more extensive perturbation caused by loss of continuous forests and continuing cooling and drying likely drove stronger disequilibrium, with butterflies following bursts of new plant lineages forming in that region. Spatial residual plots for relative phylogenetic diversity are supportive of this scenario (Figure 5), but further examination, in other herbivores, is warranted to see if such ordering effects may be more general. We hypothesize that areas with more active geologic histories will show evidence of lags in community ages between hosts and consumers/pollinators.

Phylogenetic endemism patterns for plants and butterflies as seen in CANAPE were surprisingly discordant, with much stronger plant endemism found in southern and central Mexico compared to butterflies. Mishler et al. ^25^ truncated ranges at southern edges of the region of interest, which might have led to more artificial range restrictions and higher endemism. This is particularly likely given that many widespread species with wet tropical affinities have range edges in the Yucatan of Mexico. Still, the discordance between plant and butterfly endemism is notable, and may point to fundamental differences in ecological, evolutionary, and biogeographic processes between butterflies and plants worthy of further study. We call particular attention to the strong pattern of mixed butterfly phylogenetic endemism in all of the warm deserts of North America. While more work is needed, it would be unsurprising if these warm deserts were generally areas of diversification and endemism for many clades, based on continuing floristic^74^ and faunistic work^75^.

### Importance for Conservation and Conclusions

Butterflies are under threat, perhaps represented most iconically by Monarchs and their decline^76,77^, but the entire fauna may be imperiled^78^. Results here provide a needed step for better prioritizing areas of highest conservation need. In particular, we document areas high in PE, PD, and RPD, which collectively harbor more accumulated evolutionary history^79^ and which also contain the most range-restricted lineages. These areas are likely to be high priorities for conservation^4,80^.

Within North America, there are four well-documented biodiversity hotspots: California Floristic Province, North American Coastal Plain, Madrean Pine-Oak Woodlands and Mesoamerica^81,82^. These areas were recognized due to their high plant diversity and endemism and threat of extinction^70^. While some hotspots have been long recognized, understanding where diversity is highest, rarest, and most threatened is still a work in progress; for example, the North American Coastal Plain region was only recently recognized as a hotspot^82^ and further work across the tree of life is still needed to discover areas harboring the most unique and rare diversity. Butterfly PD and PE patterns are also higher than expected within all four of these hotspots, strengthening arguments about protecting habitat in these areas. However, our results also uncovered a new hotspot showing significant endemism, relatively old lineages (based on significantly high RPD), and high PD in the warm deserts of North America. Our results make a strong case for habitat conservation across warm deserts in particular, especially because they are not already documented biodiversity hotspots.

The present study examined butterfly phylodiversity across North America and compared phylogenetic diversity metrics between butterflies and flowering plants. While the majority of phylodiversity studies conducted to date have focused on vertebrates and flowering plants, understanding the structure and past drivers of diversity in insects is critical, especially given concerns about potential rapid terrestrial declines^5^. Our work points the way to still broader future projects on insect and plant spatial phylogenetics, in order to dissect patterns at finer scales, and move beyond North America. Key next steps include further closing spatial and phylogenetic knowledge gaps globally, which would increase spatial extent and resolution beyond the coarse-scale assessment here. As well, we did not directly consider known butterfly host-plant associations in this work. This information, incorporated into a spatial phylogenetic framework, would deliver a stronger process-oriented understanding of spatial co-diversification that have shaped terrestrial ecosystems.

## Acknowledgements

We thank Map of Life team for access to range products especially for Mexican species. We also thank Caroline Storer for reviewing and providing valuable edits to the manuscript. All range products used here can be visualized at Map of Life (mol.org). CE, VB, AK, and RG were supported by the ButterflyNet project (DEB-1541500). MB was supported on a UFBI Fellowship.

## Data availability

All sequence data custom generated from this project are available on the Barcode of Life Database and will be made public upon paper acceptance. BOLD data are auto populated to Genbank on a regular schedule. Range map products can be visualized on Map Of Life (mol.org) and gridded summary products, DNA alignments, phylogenies and information about locus origin [will be] made available via a Dryad Deposition. The final phylogeny will be uploaded to Open Tree of Life (tolweb.org). Country level list data (as a .csv) for taxa with ranges outside of North America will also be made available in the same deposition. All maps presented here are available as GeoTIFFs in the dryad deposition.

