## Supplemental for "Spatial phylogenetics of butterflies in relation to environmental drivers and angiosperm diversity across North America"

**Supplemental Information 1**

### Range Map Production, Curation and Detailed Quality Control

All digitized range maps were joined into a single shapefile consisting of many spatial polygons which were clipped to only terrestrial areas within North America. We captured key attribute data for all range products including commonness and reported broad on-wing phenology. We did not digitize information about stray distributions for this work, since our key interest was range of source populations. We also therefore excluded species that stray into North America. We did not attempt in this first effort to further delineate breeding or non-breeding areas for long-distance migratory butterflies, which form only a very small percentage of species.

In order to produce highly credible range maps, we set up a rigorous review process for range map products, where maps were digitized by the authors and trained students, and a subset of the most difficult maps to digitize were reviewed by at least one member of the group to verify quality. We also instituted random spot-checks for a smaller subset of maps, especially when first starting the process. This rigor was essential because many maps included very fine-scale stippling of range locations that often required referring back to the original drawings and using a hand-lens to verify. More than a quarter of maps were carefully reviewed.

We also implemented a review process for country-level maps to assure their accuracy and to avoid issues with false positives or negatives. This curation and validation approach is as follows: 1) Country-level maps were compared to reference sources such as Wikipedia distribution descriptions and butterflies and moths of North America[1]; 2) Maps with significant disjunctions were all checked by hand. In some cases, disjunctions represented systemic issues e.g., records in French Guiana that were reported as coming from France. In other cases, disjunctions represent either real patterns or knowledge gaps across a range. Careful checking of multiple resources (e.g., Butterflies of America[2]) were used to make final validations. These country-level maps are coarse for regions outside North America, but provide a reasonable, albeit with some over-commission, approximation for range size needed for this work.

**Supplemental Information 2**

### Sequence Acquisition

**GenBank:** GenBank was searched for 13 commonly sequenced butterfly loci across all North America species using a novel toolkit that utilizes an input taxonomy list, a list of known synonyms, and a list of loci for which to search, and returns matches. Its matching approach utilizes a probe sequence for each locus of interest. In cases where there are multiple returns for a species x locus combination, the toolkit chooses the best sequence using a set of well-defined rules, often simplifying to picking the longest sequence with the most unambiguous DNA content (See https://github.com/sunray1/GeneDumper). Of the 13 loci used for this work, 12 were nuclear genes: arginine kinase (*ArgKin*), ribosomal proteins S2 and S5 (*RpS2*/*RpS5*), carbamoyl phosphate synthetase (*CAD*), catalase (*CAT*), glyceraldehyde-3-phosphate dehydrogenase (*GAPDH*), elongation factor 1 alpha (*Ef1a*), dopa decarboxylase (*DDC*), malate dehydrogenase (*MDH*), hairy cell leukemia (*HCL*), isocitrate dehydrogenase (*IDH*), and wingless (*Wgl*). Three common mitochondrial genes were also included, but were treated as one to avoid duplicate sequences: cytochrome oxidase subunits 1 (*COI*) and 2 (*COII*) and the trnL intron located between them.

Once sequences were downloaded from GenBank, they were auto-cleaned using a set of criteria built into the GeneDumper toolkit. This approach separates full-length sequences and sequence fragments. Full-length sequences are first aligned, then chunks of sequence fragments are iteratively added to this alignment using the --addfragment and --adjustdirection commands of MAFFT[3]. Since species x locus pairs frequently only have partial fragments of sequences that align to different parts of the whole locus, this approach decreases the number of misaligned fragments. Alignments were visualized using AliView v.1.26[4] and ultimately manually curated to remove flanking regions and to check that loci maintained the correct reading frame. GeneDumper uses a tiling approach for choosing sequences (in order to get the most amount of data across a locus), allowing some species x locus pairs to have slightly (0-20 bps) overlapping fragments generated from different sources. These fragments were merged together to create a 50% consensus sequence that contained data across the whole locus, where possible.

**BOLD:** Barcode of Life sequences were assembled for all species in the species list using a novel python script to query BOLD's API (http://www.boldsystems.org/index.php/resources/api) for locus information. Most of BOLD data is provisioned to Genbank but some genetic data remains unique to BOLD. If there were multiple sequences available for a species, all sequences were downloaded, aligned and a consensus sequence was created, choosing the most common nucleotide for each site. Only sequences that matched the 13 loci listed above were kept, and the vast majority of BOLD sequences are the barcode marker, cytochrome oxidase I (*COI*). Sequences were aligned using the same iterative approach described above and manually curated to remove inserts and flanking regions. We de-duplicated sequences present in both BOLD and GenBank.

**Target Capture (AHE):** We sequenced 224 samples using Anchored Hybrid Enrichment (AHE[5]) with the Butterfly1.0 target capture set of Espeland et al., (2018)[6], and the Butterfly2.0 target capture kit of Kawahara et al. (2019)[7]. Both sets include the 13 genes of interest and typically produced full-length, high quality sequences. When available, these sequences were prioritized over sequences found from other sources because their identification could be verified.

**Novel sequencing of museum specimens:** Collections in the McGuire Center for Lepidoptera & Biodiversity (MGCL), Florida Museum of Natural History, Gainesville, FL. USA, were searched for specimens for species without sequence data on public data portals. These specimens were pinned and dried, dried and papered, or stored in ethanol. One leg from each specimen was removed and transferred to a 96 well PCR microplate containing 30 uL 95% EtOH. Each well and specimen was labeled with a sample code before DNA extraction. For museum species with multiple specimens, the specimen with the most recent collection date was used to collect a tissue sample. Plates were shipped to the Canadian Centre for DNA Barcoding (CCDB) for *COI* barcoding (Sanger).

Digital voucher images of museum specimens were taken with their associated data whenever possible. For pinned specimens, we used a 12-megapixel rear-facing camera with quad-LED lights. For ethanol and papered specimens, we chose representative photos of species from Butterflies of America[1]. Photographers were contacted for reuse permission and it was noted in the submission that the photos were not of the original specimen. *COI* barcodes were received and aligned together with all other data sources using MAFFT.

**Supplemental Information 3**

**Phylogeny Reconstruction**

Cleaned and aligned sequences for each locus from each source were concatenated together and realigned. Duplicate sequences were removed and those with 95% or more similarity across overlapping regions were merged into a 50% consensus sequence using a python script. Sequences that could not be merged were submitted to BLAST[8] to determine similarity to other sequences. The sequence with the most closely related hits to the reference sequence was chosen. Once there was one sequence per species per locus, all loci were concatenated across species into a supermatrix using FASconCAT-G v.1.02[9].

Phylogenetic trees were built using maximum-likelihood phylogenetic analyses in RAxML v.8.2.10[10]. A preliminary tree was built using an unpartitioned GTR+Γ model of nucleotide evolution using a constraint tree based on family level relationships following Espeland et al., 2018[6]. That analysis recovered seven monophyletic families, including Hesperiidae and Hedylidae. Long branches (greater than 99.5% longer than average branch lengths and thus significant outliers) from this preliminary tree were removed from the alignment. Singleton GenBank and BOLD sequences with species identifiers that did not group within the correct subfamily were removed under the assumption that these represented labeling or other errors during deposition. Only 17 sequences were removed based on these criteria.

The final alignment was partitioned in three different ways: by gene, by codon and by both gene and codon. PartitionFinder v.2.1[11] was used to determine the best partitioning scheme and substitution model. Only models implemented within RAxML were searched and ranked under the AICc criterion. The best partitioning scheme was then used as input for model selection in RAxML to build 100 trees. The tree with the log likelihood score closest to 0 was chosen as the final tree.

We determined tree support by running 200 bootstraps via RAxML under the GTR+Γ+I model and partitioning scheme. Bipartitions were drawn on the final maximum-likelihood tree using two methods: Felsenstein's binary bootstrap method and a gradual "transfer" distance method implemented in BOOSTER[12]. The distance method accounts for the fact that datasets with large numbers of taxa may contain rogue taxa with unstable phylogenetic positions. Instead of removing these taxa, this method quantifies these errors as instability scores and uses these values to calculate BS scores. Phylogenies were visualized with FigTree v2.0[13] and phytools[14].

Divergence times were estimated using treePL v.1.0[15,16] using a congruification approach. We used dates from Espeland et al. (2018) as constraints on analogous nodes on the best-supported maximum likelihood tree, focusing on deeper branches of the tree. Espeland et al. (2018) describes two dated phylogenies; both phylogenies were built using nine fossil calibration points, the F48+Γ substitution model and a lognormal, birth-death tree prior, but used differing values of the median age of Angiosperms for root calibration. We used the maximum overlap between date ranges of the six families as input into treePL. Analysis options were first optimized using the ‘prime’ command. A smoothing value of 0.00001 was chosen following five cross-validation analyses. All analyses determined this value to have the lowest chi-squared value.

**Phylogenetic results and relationships**

We retrieved 21,831 sequences recovered from GenBank across all 13 loci of interest. After sequence filtering, cleaning, and the best sequence(s) for a particular species/locus pair were chosen, 3,997 sequences across 1,019 (53%) species remained. From the BOLD sequence database, 1,696 sequences were harvested across 10 of the loci of interest, representing 1,218 (63%) species. Sequencing as part of the ButterflyNet project provided 2,735 sequences across 12 of the loci of interest were found, representing 224 (11%) species and adding 11 unique species.

Subfamily and tribe-level relationships within each family generally agree with those of Espeland *et al*. (2018). Papilionidae is thought to be the sister group to the remaining butterflies and has traditionally been divided into three extant subfamilies: Baroniinae, Parnassiinae, and Papilioninae[17,18]. The sole member of monospecific family Baroniinae, Baronia brevicornis, is typically considered the sister to the other two subfamilies, but has also been grouped to Parnassiinae with Papilioninae as sister[19]. All three subfamilies are generally accepted as monophyletic, but Papilioninae has been hypothesized to be polyphyletic[6]. Our results recover Baroniinae as sister to the remainder of the family (BS = 99.5), a clade containing Parnassiinae + Papilioninae, rendering all subfamilies monophyletic (BS = 99.8 and 94.3 respectively).

Family Hesperiidae is one of the largest and most diverse families of butterflies. Although the monophyly of this group is well defined, there is disagreement on the relationships at subfamily and tribal levels[20,21]. Only four of the eight described subfamilies were sampled in our phylogeny. Three were recovered as monophyletic (Hesperiinae, Eudaminae and Heteropterinae), with Pyrginae as paraphyletic. Within Pyrginae, tribes Carcharodini, Achlyodini, Erynnini, and Pyrgini formed a single clade (BS = 97) sister to tribes Pyrrhopygini and Celaenorrhinini (BS = 89), followed by the other three subfamilies (Eudaminae + (Hesperiinae + Heteropterinae))) (BS = 97, 99 and 95, respectively).

Pieridae is currently divided into four subfamilies: Pseudopontiinae, Dismorphiinae, Coliadinae and Pierinae[22]. Pseudopontiinae is monotypic, containing a single genus found in Africa and was thus excluded from this study. Representatives from the other three subfamilies had the following relationships: (Dismorphiinae + (Coliadinae + Pierinae)), following recent studies[23,24] (BS = 100, 98 and 99). Subfamily Pierinae is usually divided into two tribes: Pierini and Anthocharidini. Our analysis supports the monophyly of these two tribes (BS = 100 and 98, respectively).

Relationships within Lycaenidae are currently unresolved and are the least well supported groups in our phylogeny (median BS = 68), but still generally follow the arrangement presented by Eliot, (1974)[25]. Eliot classified Lycaenidae into seven subfamilies, of which we have representatives from four (Miletinae, Theclinae, Polyommatinae and Lycaeninae). Like

Espeland et al. (2018), we find Polyommatinae were nested within Theclinae. We also find Lycaeninae nested within Theclinae. The classification presented by Corbet et al. (1992)[26] reduced Theclinae, Polyommatinae, and Lycaeninae to an inclusive “Lycaeninae”. Our phylogeny agrees with this lumping (BS = 99). Miletinae, although represented by only one species, was placed sister to these three subfamilies (BS = 100).

Riodinidae are mostly found in the Neotropics and are divided into three subfamilies: Riodininae, Euselasiinae and Nemeobiinae[27]. Nemeobiinae are strictly Old World riodinids and were therefore excluded from this study. The two remaining subfamilies, Riodininae and Euselasiinae, were both monophyletic with 100 and 98 BS support, respectively. Tribal relationships within Riodininae were still mixed and unsupported in all studies. This may be due to rapid radiations within this subfamily[28]. Although there was some paraphyly between tribes, we agree with the suggestion made by Espeland et al. (2018) that an Emesis-Apodemia tribal group might need to be erected (BS = 94).

There is no phylogenetic consensus within Nymphalidae, the most speciose butterfly family, although recent studies have shed some light on its structure[29,30]. Due to the sheer number of species (>6,000) and high amounts of diversity, Nymphalidae has been split into anywhere from 9 to 12 families[31–33]. All subfamilies in our analyses were monophyletic. We found Libytheinae to be sister to Danainae and Ithomiinae (lumped into Danainae by some) (BS = 98), which is in turn sister to the rest of the family (BS = 96). This is also one of the topologies suggested by Espeland et al. (2018). Placement of the remaining nymphalids was largely concordant with the relationships in Espeland et al. (2018), with the exception of the placement of Apaturinae. Espeland et al. (2018) found Apaturinae to be sister to Biblidinae whereas we found Apaturinae sister to the clade containing Cyrestinae and Nymphalinae. This may be due to the lack of representation of subfamily Pseudergolinae in this study since it is found in Asia, not North America.

### Supplemental Information References


1. Lotts, K. & Naberhaus, T. Butterflies and Moths of North America. https://www.butterfliesandmoths.org/ (2017) doi:http://www.butterfliesandmoths.org/.
2. Warren, A. D. *et al.* Illustrated Lists of American Butterflies. http://www.butterfliesofamerica.com/ (2016).
3. Katoh, K. & Standley, D. M. MAFFT: Iterative refinement and additional methods. *Methods in Molecular Biology* **1079**, 131–146 (2014).
4. Larsson, A. AliView: a fast and lightweight alignment viewer and editor for large datasets. *Bioinformatics* **30**, 3276–3278 (2014).
5. Lemmon, A. R., Emme, S. A. & Lemmon, E. M. Anchored Hybrid Enrichment for Massively High-Throughput Phylogenomics. *Systematic Biology* **61**, 727–744 (2012).
6. Espeland, M. *et al.* A Comprehensive and Dated Phylogenomic Analysis of Butterflies. *Current Biology* **28**, 770-778.e5 (2018).
7. Kawahara, A. Y. *et al.* Phylogenetics of moth-like butterflies (Papilionoidea: Hedylidae) based on a new 13-locus target capture probe set. *Molecular Phylogenetics and Evolution* **127**, 600–605 (2018).
8. Altschul, S. F. *et al.* Gapped BLAST and PSI-BLAST: A new generation of protein database search programs. *Nucleic Acids Research* vol. 25 3389–3402 (1997).
9. Kück, P. & Longo, G. C. FASconCAT-G: Extensive functions for multiple sequence alignment preparations concerning phylogenetic studies. *Frontiers in Zoology* **11**, (2014).
10. Stamatakis, A. RAxML version 8: a tool for phylogenetic analysis and post-analysis of large phylogenies. *Bioinformatics* **30**, 1312–1313 (2014).
11. Lanfear, R., Frandsen, P. B., Wright, A. M., Senfeld, T. & Calcott, B. PartitionFinder 2: New Methods for Selecting Partitioned Models of Evolution for Molecular and Morphological Phylogenetic Analyses. *Molecular Biology and Evolution* msw260 (2016) doi:10.1093/molbev/msw260.
12. Lemoine, F. *et al.* Renewing Felsenstein's phylogenetic bootstrap in the era of big data. *Nature* **556**, 452–456 (2018).
13. Rambaut, A. FigTree v2.0. *Institute of Evolutionary Biology, University of Edinburgh, Edinburgh.* (2010).
14. Revell, L. J. phytools: an R package for phylogenetic comparative biology (and other things). *Methods in Ecology and Evolution* **3**, 217–223 (2012).
15. Sanderson, M. J. Estimating Absolute Rates of Molecular Evolution and Divergence Times: A Penalized Likelihood Approach. *Molecular Biology and Evolution* **19**, 101–109 (2002).
16. Smith, S. A. & O'Meara, B. C. treePL: divergence time estimation using penalized likelihood for large phylogenies. *Bioinformatics* **28**, 2689–2690 (2012).
17. Allio, R. *et al.* Whole Genome Shotgun Phylogenomics Resolves the Pattern and Timing of Swallowtail Butterfly Evolution. *Systematic Biology* **69**, 38–60 (2020).
18. Condamine, F. L., Nabholz, B., Clamens, A. L., Dupuis, J. R. & Sperling, F. A. H. Mitochondrial phylogenomics, the origin of swallowtail butterflies, and the impact of the number of clocks in Bayesian molecular dating. *Systematic Entomology* **43**, 460–480 (2018).
19. Nazari, V., Zakharov, E. v. & Sperling, F. A. H. Phylogeny, historical biogeography, and taxonomic ranking of Parnassiinae (Lepidoptera, Papilionidae) based on morphology and seven genes. *Molecular Phylogenetics and Evolution* **42**, 131–156 (2007).
20. Yuan, X., Gao, K., Yuan, F., Wang, P. & Zhang, Y. Phylogenetic relationships of subfamilies in the family Hesperiidae (Lepidoptera: Hesperioidea) from China. *Scientific Reports* **5**, (2015).
21. Zhang, J., Cong, Q., Shen, J., Brockmann, E. & Grishin, N. v. Genomes reveal drastic and recurrent phenotypic divergence in firetip skipper butterflies (Hesperiidae: Pyrrhopyginae). *Proceedings of the Royal Society B: Biological Sciences* **286**, 20190609 (2019).

22. Braby, M. F., Vila, R. & Pierce, N. E. Molecular phylogeny and systematics of the Pieridae (Lepidoptera: Papilionoidea): Higher classification and biogeography. *Zoological Journal of the Linnean Society* **147**, 239–275 (2006).
23. Cao, Y., Hao, J. S., Sun, X. Y., Zheng, B. & Yang, Q. Molecular phylogenetic and dating analysis of pierid butterfly species using complete mitochondrial genomes. *Genetics and Molecular Research* **15**, (2016).
24. Ding, C. & Zhang, Y. Phylogenetic relationships of Pieridae (Lepidoptera: Papilionoidea) in China based on seven gene fragments. *Entomological Science* **20**, 15–23 (2017).
25. Eliot, J. N. The higher classification of the Lycaenidae (Lepidoptera) : a tentative arrangement. *Bull Br Mus Nat Hist Entomol* **28**, 371–505 (1974).
26. Corbet, A. S., Pendlebury, H. M. & Eliot, J. N. *The butterflies of the Malay Peninsula.* (Malayan Nature Society, 1992).
27. Seraphim, N. *et al.* Molecular phylogeny and higher systematics of the metalmark butterflies (Lepidoptera: Riodinidae). *Systematic Entomology* **43**, 407–425 (2018).
28. Espeland, M. *et al.* Ancient Neotropical origin and recent recolonisation: Phylogeny, biogeography and diversification of the Riodinidae (Lepidoptera: Papilionoidea). *Molecular Phylogenetics and Evolution* **93**, 296–306 (2015).
29. Chazot, N. *et al.* Priors and Posteriors in Bayesian Timing of Divergence Analyses: The Age of Butterflies Revisited. *Systematic Biology* **68**, 797–813 (2019).
30. Wahlberg, N. *et al.* Nymphalid butterflies diversify following near demise at the Cretaceous/Tertiary boundary. *Proceedings of the Royal Society B: Biological Sciences* **276**, 4295–4302 (2009).
31. Brower, A. V. Z. Phylogenetic relationships among the Nymphalidae (Lepidoptera) inferred from partial sequences of the *wingless* gene. *Proceedings of the Royal Society of London. Series B: Biological Sciences* **267**, 1201–1211 (2000).
32. Freitas, A. V. L. & Brown, K. S. Phylogeny of the Nymphalidae (Lepidoptera). *Systematic Biology* **53**, 363–383 (2004).
33. Wahlberg, N., Weingartner, E. & Nylin, S. Towards a better understanding of the higher systematics of Nymphalidae (Lepidoptera: Papilionoidea). *Molecular Phylogenetics and Evolution* **28**, 473–484 (2003).

### Supplemental Figure 1

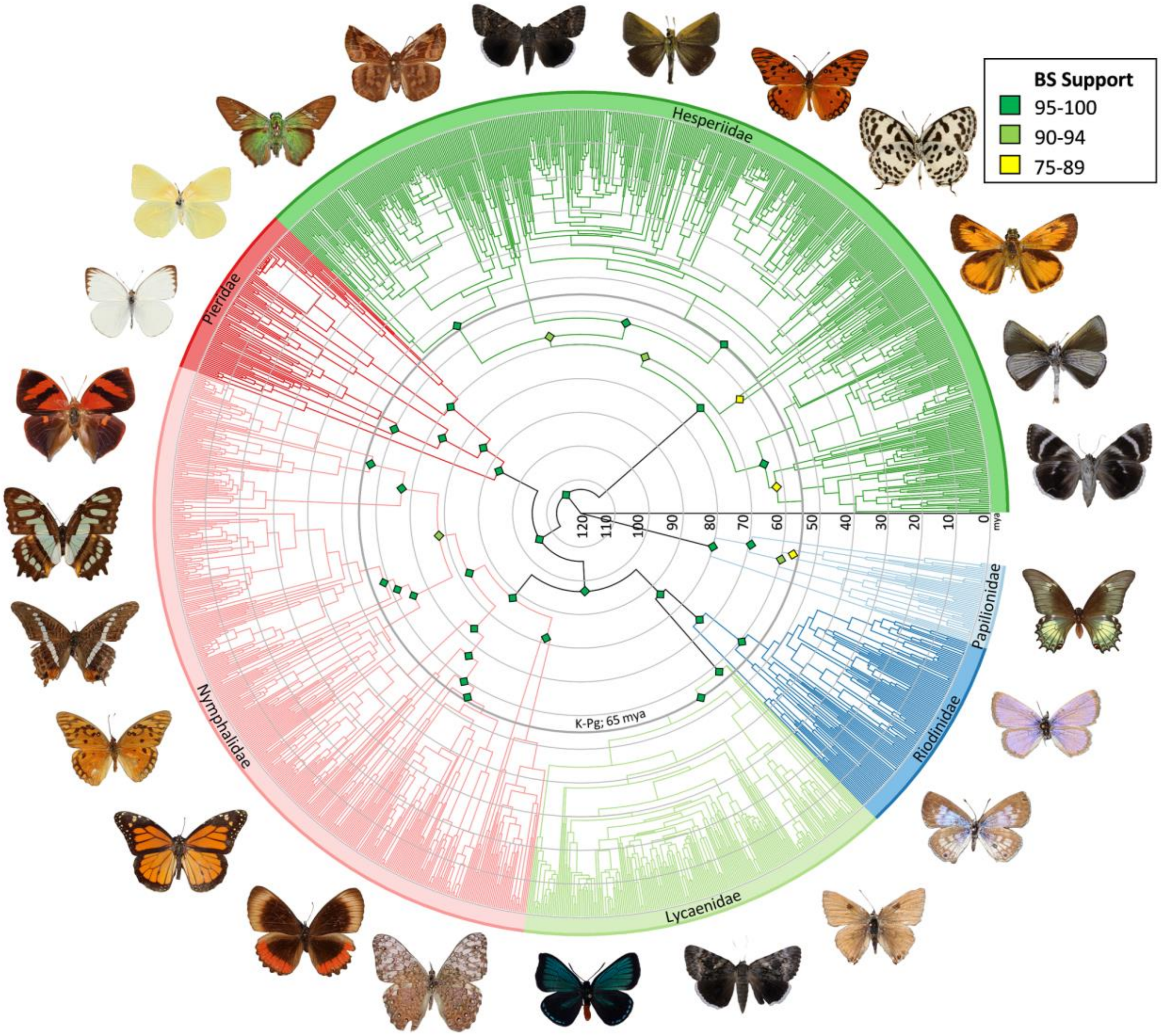

**Supplemental Table 1**

The length of the 14 loci used as the starting sequence for input into GeneDumper.

| Locus Name | Locus Length |
|---|---|
| *ArgKin* | 596 |
| *CAD* | 2,211 |
| *CAT* | 1,293 |
| *COI_trnL_COII* | 2,271 (*COI*: 1,531; *trnL*: 67; *COII*: 673) |
| *DDC* | 957 |
| *EF1a* | 1,240 |
| *GAPDH* | 609 |
| *HCL* | 633 |
| *IDH* | 709 |
| *MDH* | 733 |
| *RpS2* | 474 |
| *RpS5* | 614 |
| *Wgl* | 453 |

**Supplemental Table 2**

Ages of all families (in millions of years) in this study and others with 95% confidence intervals.

| Clade | Wahlberg et al. (2009) | Heikkilä et al. (2012) | Espeland et al. (2018) * | Condamine et al. (2018) | Chazot et al. (2019) | This study |
|---|---|---|---|---|---|---|
| Papilionoidea | 104 (93-116) | 110 (92-128) | 119 (91-143) | 98 (66 - 189) | 108 (89-129) | 119.50 |
| Papilionidae | 63 (93 - 116) | 75 (62-88) | 84 (63-109) | 86 (56-164) | 68 (53-84) | 79.67 |
| Hesperiidae | N/A | 65 (54-79) | 79 (60-99) | 76 (55-143) | 65 (56-78) | 72.49 |
| Pieridae | 73 (57-86) | 80 (67-97) | 87 (67-108) | 71 (45-135) | 77 (63-92) | 92.29 |
| Lycaenidae | 75 (63-86) | 73 (58-84) | 78 (60-96) | 62 (39-119) | 71 (57-85) | 57.40 |
| Riodinidae | 65 (56-76) | 72 (57-83) | 73 (56-92) | 60 (36-116) | 73 (60-88) | 72.49 |
| Nymphalidae | 94 (84-104) | 87 (74-101) | 91 (71-112) | 80 (52-151) | 82 (68-98) | 87.43 |

* Dates used in Espeland et al. (2018) were used to constrain the tree in the present study.
